# Modeling assembly dynamics and stability of microbial communities

**DOI:** 10.64898/2025.12.17.694811

**Authors:** Andreas Eilersen, Kim Sneppen, Sebastian Bonhoeffer

## Abstract

Predicting the species composition of microbial communities is a problem that has proven surprisingly difficult, including in fully controlled microbial systems in a lab setting. Few organisms are able to establish themselves in an community with no others, even if nutrients are provided. At the same time some species are observed to be unable to stably coexist with certain other species. We here present a model attempting to reduce the dynamics of community assembly to its most basic components, while preserving its salient features. The model deals not with abundances or densities of bacterial populations or resources, but only with their presence or absence. Similarly, only three types of discrete relationships between species are considered: species excluding each other, and mutualistic dependence on and production of nutrients in the form of exometabolites. Despite these simplifications, the model system still exhibits rich dynamics, including extinction cascades and emerging stability against invasion. We derive conditions for these to occur and compare our results with existing knowledge of microbial communities. Our results provide a novel approach for the theoretical study of microbial communities and their stability.

**Significance:** Understanding the processes governing the community composition of microbial ecosystems is an as-yet unsolved problem. In this article, we propose a novel rule-based model for the assembly and stability of communities of microbial species. Despite its simplicity the model exhibits rich dynamics, including critical points and extinction cascades. It presupposes no knowledge of specific population parameters or exact resource or species abundances, but focuses on pairwise interactions between species and their metabolites. The framework may be useful for understanding how real microbial communities arise and what causes them to break down.

---

**T**he assembly and collapse of ecosystems has received increasing attention in the last few decades. This has long been the case within ecology and paleobiology, where a model for understanding ecosystem stability and mass extinction events has long been sought (1, 2). More recently bacterial communities and microbiomes, particularly in the context of the human gut, have similarly raised questions about the principles of microbiome assembly and stability (3–5). It has turned out that the symbiosis with the various microbiomes of the body has a range of effects on the host, which may be more profound and varied than previously thought (6–8). Increasing our understanding of what makes an ecosystem (un)stable might thus not only have implications for ecology and biodiversity but also for human health.

Along these lines, some experimental and observational work has been done on ecosystem assembly, attempting to devise rules for putting together microbial communities. This has taught us that some pairs of microbial species coexist readily, while others are seemingly unable to coexist (5). In such exclusionary pairs, the introduction of one species leads to the extinction of the other. This raises the question whether such seemingly simple “all-or-nothing” rules for assembly of microbial communities can be used to create more tractable models for microbial community structure and diversity.

Additional studies have shown that stochastic effects may play a large role on gut microbiome assembly (9, 10), and a study in fruit flies has shown that previous colonization by a highly competitive bacterial species makes the gut stable to invasion by less competitive later invaders (11). Generally, priority effects seem to affect the final composition of microbiomes (12). Taking a broader view of the entire network of coexistences and interactions, it has been shown that bacterial co-occurrence networks are often highly inhomogeneous, with some bacteria co-occurring with many others, but most having relatively few such associations. Microbial communities are known to be organised hierarchically along trophic lines (13), and in non-microbial communities, interaction networks are similarly strongly asymmetric (14) and nested (15). Concretely, these observations imply that interactions usually have a clear direction (species A affects B, but often not the other way around), and that specialist species interact with generalists at an above-chance rate, but rarely with other specialists.

Inspired by these discoveries, we here develop a simple computational model for species coexistence and competition and use it to make predictions about ecosystem stability and extinction events. We use the available knowledge about microbiome structure and food networks to inform our assumptions. Thereby, we hope to provide a plausible theoretical framework for understanding among other things the dynamics behind extinction events, and the requirements for ecosystem stability against extinctions and invasions.

Some similar, minimalist and often physics-inspired models of ecosystem assembly and species extinction have been proposed before (16). One notable example is the Bak-Sneppen model, which proposes that the distribution of extinction events is due to self-organised criticality (17). Other models explicitly consider the population dynamics of invaders and residents (16, 18). This has the advantage of being more biologically realistic and enabling the modelers to use the significant existing body of knowledge of Lotka-Volterra dynamics. However, although some research on interaction parameters in human microbiomes has been done (19) the parameterization of the models is difficult to do in a realistic manner, and their dynamics may end up being too complex for a straightforward analysis. Furthermore, some experimental evidence from the study of microbiomes specifically indicates that it is the presence or absence - rather than the abundance - of a given species within the microbiome which shapes host traits (20).

Here, we will attempt to do away with the specifics of species interactions and abundances by not modeling the population dynamics of species. Instead, we will view their presence or absence as a binary variable. A similar approach was taken by Clegg & Gross (21), who looked at the feasibility of communities of microbes cross-feeding on each other’s metabolites. One main difference between their work and ours is that where their work focuses on the instantaneous feasibility of a certain community given a cross-feeding network, we will focus on assembly dynamics over time. Analyzing temporal assembly and invasion dynamics will require a different analytical approach. A further novelty of our study is that it, in addition to cross-feeding, will attempt to describe the effect of competition and antagonism. Finally, we will test our model with two types of cross-feeding networks, namely one random (Erdős-Renyi) and one hierarchical “trophic” network.

We will assume that competing species are antagonistic to the point of total exclusion. Nutrients follow equally simple rules: if a species in the ecosystem produces a nutrient, the nutrient is simply present, regardless of concentration. If no species produces it and it is not a basic, naturally occurring resource, it is simply absent. A bacterial species may need a nutrient, meaning that it is unable to exist if said nutrient is not available. The idea is that this will enable us to more easily describe the evolution and statistical mechanics of the community as a whole. While we have no illusion that this will capture the intricacies of microbial species interactions, we hope to better understand the basic factors shaping the composition and long-term stability of microbiomes. In addition, it is our hope that other researchers working with the assembly and correlations of microbial communities will find this style of description useful and apply it as a tool for their concrete work on specific communities.

## Model

To construct a model of ecosystem assembly and biodiversity, we incorporate two basic interactions between species: competition leading to exclusion, and symbiosis. Symbiosis is mediated by a limited set of nutrients. Nutrients are provided either naturally in the environment (“basic” nutrients) or through exometabolite production by bacterial species. Bacteria produce some number *n*_*pro*_ of nutrients, and will in turn require a number *n*_*req*_ of other nutrients to exist. Bacterial crossfeeding and symbiosis thus take the form of a bipartite network with links from bacteria to nutrients, and nutrients to bacteria, but not directly between bacteria.

Besides this, antagonistic interactions between species are modelled as an additional directed network of negative links directly between species. The negative interaction network is thus a directed graph, but as opposed to the symbiotic interaction network, it is not bipartite. If there is a negative link from species A to species B, and A is present, B is unable to persist in the system (but not vice-versa). A diagram illustrating the network underlying the model can be seen in fig. 1.

**Fig. 1.**
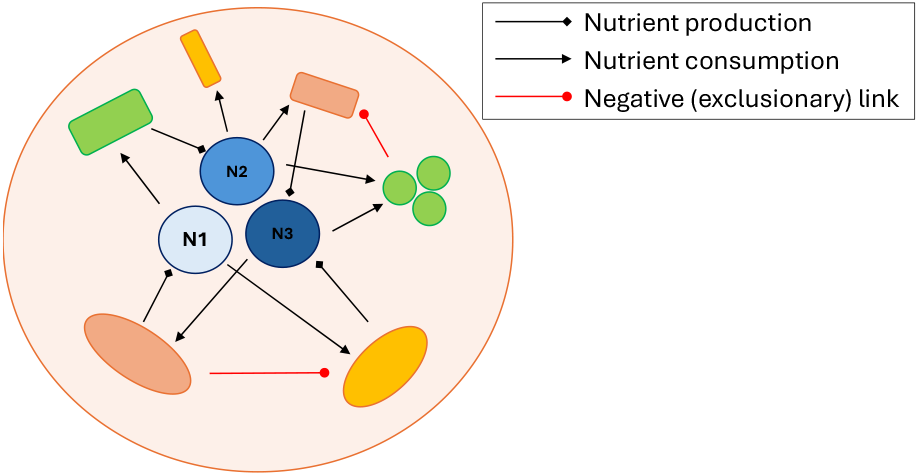
A schematic illustration of the model, showing a network of bacterial species (colored shapes) and their exometabolites (blue, numbered circles). Some of the species support the existence of others through nutrient production (black arrows) and some exclude others (red, blunt arrows).

To simulate the temporal invasion dynamics, we take one species per timestep and let it attempt to invade. If the system contains all necessary nutrients and there are no antagonists present, the invasion is successful. Then, we check if the new invader excludes any species already present. If so, they go extinct. Finally, we check if these extinctions have led to any nutrients no longer being produced. If so, the species depending on the depleted nutrients also go extinct, which may in turn lead to more nutrient depletion and more extinctions - what we refer to as a cascade. We therefore iterate the last step until no more nutrient depletions happen.

### Network structure assumptions

For the analytical derivations given below, we will assume that the negative links between species are randomly distributed, and the number of links per species is thus given by a binomial distribution. The same holds for the number of nutrients produced and required by each species. The negative interaction network is thus a (monopartite) Erdős-Renyi network, and the nutrient production and consumption network is a random, bipartite graph. We will retain this assumption for half of our numerical treatment. After this, we will attempt to move beyond it, since many natural interaction networks between species (microbial or otherwise) appear to be scale-free (22), nested (15), and hierarchical, separated into trophic levels (13). This would indicate that at least symbiotic relationships between species by way of nutrient production should follow a much more skewed distribution, with most species having few symbionts dependent on their exometabolites, but a few species being essential for a very large number of symbionts.

In the second half of our numerical treatment, we will therefore use what is arguably a more naturalistic set of networks. To generate it, we will assume that naturally occurring microbiomes are founded upon a supply of highly complex “raw” nutrients. This is in keeping with the main points of Gralka *et al*.(13). The complex nutrients could be bits of plant matter, organic soil particles or similar. Relatively few species can break down and metabolize each of these. The result of this primary metabolism is a set of less complex nutrients allowing for more “picky” species to survive, which further break down the nutrients to ever more simple compounds. Finally, we end up with simple organic molecules such as glucose and lactose which are very widely consumed.

We will construct this trophic model network by letting a number of nutrients be freely available, but necessary only for a small number of species each. These species in turn produce a smaller set of nutrients, consumed by a larger set of species, and so on. The network is generated such that the degree distribution giving the number of outgoing links from species to species (mediated by nutrients) is a decreasing function of degree. That is, most species do not support the existence of that many others, while a small number of “degrader” species support a large number of “consumers”. The negative interaction network will still be random. In both network types described above, there will be no correlation between the number of antagonistic and the number of symbiotic links. In the random-network case, the out-degree of each species (how many nutrients it produces) will also be unrelated to its in-degree (how many it consumes), whereas in the trophic network case, both of these will be related to position in the trophic hierarchy.

Having constructed these networks, we will investigate whether these combinations of networks behave differently when looking at the probability of collapse or extinction cascades, and at the stability against invasion.

### Analytical results

It is worth considering what can be said about the model dynamics from a mathematical analysis of its basic assumptions. For the purposes of the following, we will only work with the case where all network links are randomly distributed.

An important question to ask before analysing these models is “what do we mean by stability?”. We may gain some intuition from looking at real microbial communities. They seem to maintain stability by (1) having functional redundancy, i.e., several species producing every essential nutrient, and (2) excluding potentially harmful species from invading (23). When such invasions happen, they may suffer an overgrowth of the invader, potentially resulting in collapse. The first part of this statement points towards the probability of nutrient depletion and extinction cascades as the relevant measure of stability. At the same time, the other part of the statement instead suggests invasibility as such a measure. If an invader can trigger a cascade, it may pave the way for itself to take over the system. These suggested measures correspond roughly to two widely used ecological stability measures, namely community resilience (the ability to recover the initial state after a perturbation) and resistance (how hard it is for an invader to establish itself) (24). Therefore, these are the definitions of stability that we will use in the following analyses of the model. Numerical verification of the results of this section can be found in the supplementary information.

#### Equilibrium diversity

The number of excluded species is directly proportional to the number of resident species, which we will here call *x*. For the system to be stable against invasion, every non-resident species must be excluded by at least one resident species. On average, the number of species excluded by those present will be *Np*_*neg*_*x*, where *p*_*neg*_ is the probability that any given negative link from one species to another exists, and *N* is the total number of available species in the environment. Of these, a fraction (1 − *x/N* ) will be non-resident species. For stability against invasion to be likely, the average number of excluded species should be equal to the number of non-resident species:

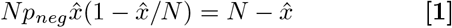

where 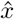 is the number of species present in the non-invasible system. Solving this quadratic equation, we get

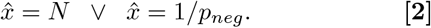

For the system to be in equilibrium, i.e., the number of invasions being equal to the number of extinctions, the expected number of invasions per timestep must be equal to the expected number of extinctions per timestep. Since the timestep is defined as the time between invasion attempts, the expected number of invasions will be the probability of success, *p*_*inv*_. In turn, the expected number of extinctions per timestep must be *p*_*inv*_*p*_*neg*_*x*, since any successful invader will exclude *p*_*neg*_*x* species on average. For equilibrium, we thus have *p*_*inv*_*p*_*neg*_*x*^*^ = *p*_*inv*_, implying

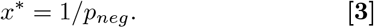

Thus, whether our stability condition is constant diversity (*x*^*^) or resistance to invasion (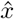), we get that the number of species present depend on the number of negative links as *x* = 1*/p*_*neg*_.

#### Resilience

We predict that a prominent feature of this model will be extinction cascades resulting from one extinction leading to depletion of a nutrient, leading to more extinctions. The risk of this happening will be inversely related to the resilience of the system. We can calculate the probability of depletion of at least one nutrient as follows:

In a community of *x* species, the probability that a nutrient is not produced is (1 − *p pro* )*x*, where *p pro* is the probability of any given positive link from bacterium to nutrient existing. *p*_*pro*_ is related to the number of nutrients produced per species, *n*_*pro*_, as *p*_*pro*_ = *n*_*pro*_*/N*_*nut*_ (this is at least the case if the links are randomly distributed). From this, we get the probability that all nutrients are produced as

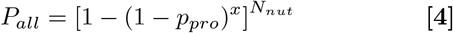

At equilibrium, the substitution *x* → 1*/p neg* can be made, since if all nutrients are produced, the only factor limiting *x* is exclusion by antagonistic species. If we define a cascade as starting with a nutrient being depleted, the probability of cascades is thus 1 − *P all*. It is clear from these equations that the probability of cascades falls rapidly with increasing diversity *x*. Therefore, if *x* ≈ 1*/p neg* is much greater than the mean number of species required to maintain all nutrients, *N*_*nut*_*/n*_*pro*_ = 1*/p*_*pro*_, cascades are highly unlikely. Cascades are thus mostly expected to be a phenomenon occurring near this critical number of species. The expression for *P*_*all*_ derived here nonetheless only gives an indication of the threshold for cascades, since in the model multiple species can go extinct at once, reducing stability. On the other hand, the assembly rules ensure that the least picky species become residents first, resulting in greater stability.

#### Resistance to invasion

Having shown that the number of negative links between species is the most important parameter for the end state of the system, we can now explore how it concretely affects the dynamics. The probability that an invasion cannot happen at all is given by the expression

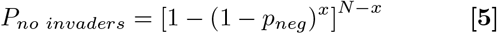

derived by negating the probability that any non-resident species can invade, in turn itself a negation of the risk of a random invading species being excluded by a resident. Substituting the result derived above, that *x* ≈ 1*/p*_*neg*_ at equilibrium, we get

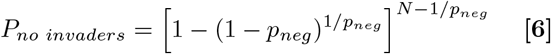

This function depends very strongly on *p*_*neg*_ and for *p*_*neg*_ *<< N* only weakly on *N* . It also drops to a very small value as *p*_*neg*_ approaches 1*/N* . With this in mind, we can say with some confidence that the system will quickly lose its stability when *p*_*neg*_ *>* 1*/N*, regardless of the system size. Alternatively, it can be said that the system becomes unstable if the average number of negative links per species, *n*_*neg*_ = *Np*_*neg*_ is greater than 1.

As above, which species are resident and which are not is of course not randomly chosen, but rather subject to a sorting effect, as highly competitive species (with many outgoing negative links) are likely to become residents quickly, whereas those with many ingoing negative links are likely to remain excluded. Therefore, it is likely that stability can be maintained for *n*_*neg*_ slightly greater than 1. Unfortunately, we have not been able to estimate the effect of this analytically.

#### Summary and interpretation

Perhaps the most important take-away of these results is the role of *p*_*neg*_ in limiting the number of species that can stably coexist in a community, as given in eqs. (2,3). From this result, we predict there to be an inverse relation between the diversity of a community and the propensity of its potential constituent species to have strongly antagonistic interactions with other potential constituents. If a microbial ecosystem draws its members from a pool of species with many interactions that are antagonistic to the point of exclusion, we would therefore expect this ecosystem to exhibit low diversity, and vice versa.

With regard to nutrients and cross-feeding, the results are less clear. From eq. (4), we predict that the probability function of all species having their nutrient requirements met will exhibit a strong switch-like behaviour. If the number of nutrients involved in the total metabolism of the microbial community is significantly larger than the number of species, we predict the system to be unstable, with mass extinctions likely. On the other hand, if the number of species is larger than the number of nutrients, the probability that every nutrient will have some producer and some consumer quickly grows. This dynamic is thus mostly important in practice near the critical point where there exists one species for each nutrient. Here, the community will exist and have a significant diversity, but it will be unstable. Notably though, this prediction does not take into account a possible trophic structure of the microbe-resource network.

Finally, from eq. (6), we predict low-diversity communities existing in an environment with many potentially invading species to be unstable. Such small communities in rich environments will be overwhelmingly likely to experience repeated invasions, destabilising them. This holds even in the case where resident species have many exclusionary links to non-resident species, since it is likely that at least a few of them will be able to invade anyway and in turn displace some residents themselves. In the discussion, we will outline some scenarios likely to occur naturally that could be used to test these predictions.

### Numerical results

#### Random network

Numerical methods allow us to investigate the stability of the community in additional ways. One such way is the distribution of extinction cascades as a function of their size. An overview of these distributions when varying the different model parameters can be seen in fig. 2. From these plots, we see that the cascade size distributions resemble shallow exponential functions, mostly with a steeper exponential cutoff at large sizes.

**Fig. 2.**
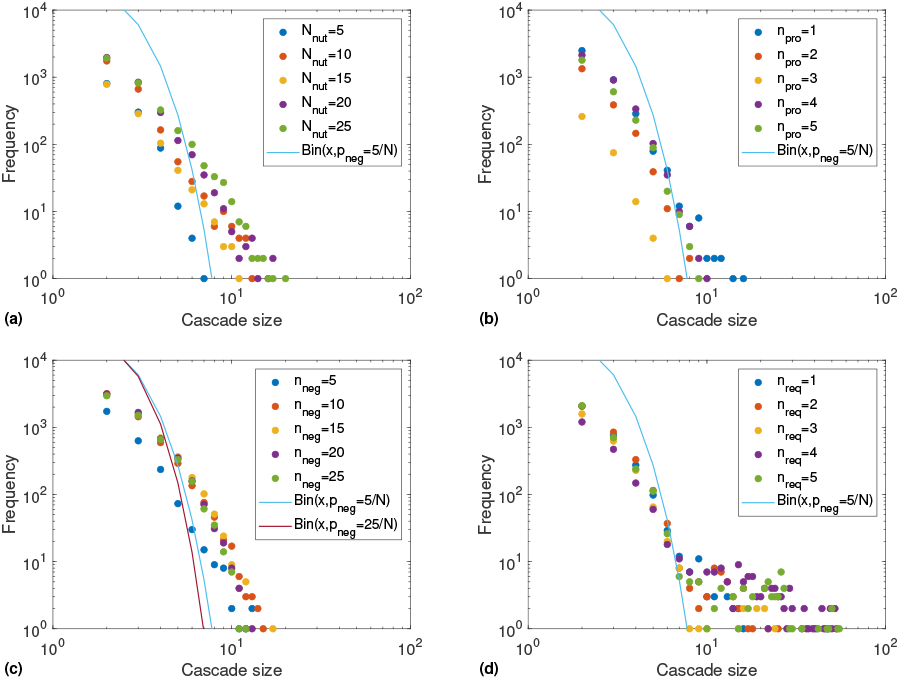
Frequency distributions of cascades by size (i.e., number of extinctions). (a) Varying the number of potentially available nutrients, *Nnut*. (b) Varying the number of nutrients produced per species, *npro*. (c) Varying number of negative links per species, *n*_*neg*_ . (d) Varying number of necessary nutrients per species, *n*_*req*_ . The parameters used are *n*_*neg*_ = 5, *N*_*nut*_ = 10, *n*_*req*_ = 1, *n*_*pro*_ = 1, and *N* = 200. For comparison, we show the cascade distributions if cascade sizes were drawn from a binomial distribution with success chance *p*_*neg*_ and *x* = 1*/p*_*neg*_ attempts. Simulations are run for 10^5^ timesteps.

The effect of changing the different parameters varies. Increasing the number of nutrients in the system, *N*_*nut*_ almost monotonically increases cascade frequency and size but does not change the shape of the distribution. On the other hand, the number of nutrients produced per species, *n*_*pro*_, slightly decreases cascade frequency and size as it grows from 1 to 3, and subsequently increases it again. The reverse is true for the number of negative links, *n*_*neg*_. The number of nutrients required per species, *n*_*req*_, is the only parameter that changes the shape of the distribution: when *n*_*req*_ is larger, the tail of the distribution becomes longer, that is, very large cascades become possible. In all cases, the slope of the distribution appears to be around − 3, though with some curvature suggesting an exponential rather than a power law distribution. Thus, large cascades are quite unlikely in any case, and as the cutoff is usually located just above a cascade size of ten extinctions, this seems to be the maximum size of cascades. We do assume that the number of species is significantly larger than the number of nutrients, but even having the two numbers be of the same order does not greatly increase the size of cascades (see supplementary).

To interpret this in words, the more nutrients are involved in the system, the more possibilities there are for initiating a cascade. One could say that a large ratio of nutrients to species decreases redundancy. This leads to more and to some extent larger cascades. The number of nutrients produced per species and the number of negative links have unclear relationships with the number of cascades. Particularly, *n*_*neg*_ reduces the diversity of the system, leading for all else being equal to smaller cascades. On the other hand, it also destabilises the system, as we shall see below. Finally, the number of nutrients that each species is dependent upon, *n*_*req*_, increases the likelihood of large cascades. It can be said to increase the connectivity of the symbiotic network between bacteria, meaning that the consequences of an extinction affects more species.

Of course, given the steeply declining size distribution function, we might guess that the measured cascades are simply invasions by highly competitive species, driving many others to extinction. However, given that we assume the network of negative interactions to be an Erdős-Renyi network, the size of such extinction events would be binomially distributed like the number of links per node in such a network. In fig. 2, we therefore plot the expected cascade size distribution if cascade sizes were binomially distributed for comparison. We observe that the binomial distribution does not fit the observed cascade distributions very well - for example, it tends to drop off more quickly at larger cascade sizes and predicts a higher number of single-extinction events.

A different measure of stability would be the invasibility of the system, which is also the opposite of the above-mentioned resistance. Therefore, we vary the four main parameters and measure the mean time between successful invasions. In addition, we record the time it takes for the system to settle down to a stable state, which we here term the settling time, measured as the time until the last successful invasion. The resulting plots are shown in fig. 3.

**Fig. 3.**
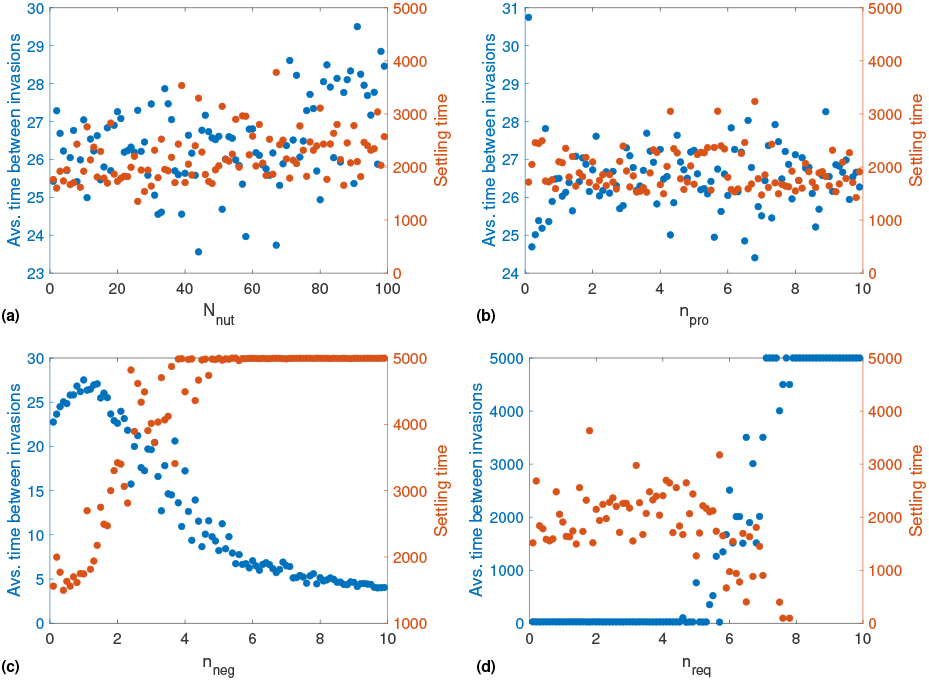
The stability of the community against invasion, measured by mean time between successful invasion attempts (left y-axes) and time to last successful invasion (right y-axis), shown as functions of (a) *N*_*nut*_, (b) *n*_*pro*_, (c) *n*_*neg*_, and (d) *n*_*req*_ . Note that the scales of the left y-axes are different. The displayed data are means of five runs - hence, when it is random whether the system reaches stability at all or not, this tends to lead to settling time values around *T/*2. Only varying *n*_*neg*_ and *n*_*req*_ seem to have a significant and consistent effect, whereas it seems to be random whether the system ends up in a stable configuration in the other cases. The parameter values are *N* = 200, *N*_*nut*_ = 10, *n*_*neg*_ = 1, *n*_*pro*_ = 1, and *n*_*req*_ = 1.

As predicted analytically, it turns out that increasing the number of negative links per species has the largest effect on stability. At *n*_*neg*_ below about 3 per species, the system will relatively quickly settle down to a stable, non-invasible state. With more negative links, however, such a stable state becomes increasingly unlikely. This is a slightly higher threshold than expected; analytically, we would expect instability to grow quickly once there is one or more negative links per species on average.

Interestingly, as *p*_*neg*_ increases and the final diversity of the system thus decreases, mean time between successful invasions initially increases slightly as well, indicating greater stability. This is likely due to a combination of two factors. Firstly, any potential invader will be more likely to be excluded, meaning that there will be more unsuccessful invasion attempts. Also, the final diversity will be slightly smaller, making complete assembly to stability quicker. What this also indicates is that the measure of stability used here is not completely trivial. Rather than showing increasing stability as we approach the trivially stable case where all species are present (and therefore, no-one can invade), we see that some mostly, but not completely, full ecosystem actually maximizes stability. The value of *p*_*neg*_ that results in maximal stability is large enough to ensure that some species are unable to invade, but small enough that established species are unlikely to be displaced. The importance of negative interactions in stabilizing a microbial community has been described by (25), who nevertheless use a different methodology and therefore arrive at a much higher threshold for system destabilization.

As with *n*_*neg*_, the nutrient requirement parameter *n*_*req*_ does appear to have a major effect on resistance looking at the plot in fig. 3 alone. Looking at the system during the simulations, the apparently highly stable states at high *n*_*req*_ turns out to be the system becoming unable to assemble at all when species are highly demanding. The stable states are therefore actually empty systems.

#### Alternative network structures

To investigate the effect of different network structures, we generate a hierarchically organised nutrient-bacteria network mimicking the trophic networks common in natural microbial communities (13). Despite the different structure, the change of network has a small effect on the resistance, for which the plots resemble those of fig. 3. One exception is the invasibility as a function of available nutrients *N*_*nut*_, which is much more variable than in the random network case. The resource production parameter *n*_*pro*_ exhibits a similar effect in both network types. Number of required nutrients per species, *n*_*req*_, is not meaningful for the trophic model, since this varies with position in the trophic hierarcy. Plots of the invasibility and settling times as functions of *n*_*neg*_ and *N*_*nut*_, and of the corresponding cascade size distributions are shown in fig. 4 for comparison. On the other hand, the slope of the cascade size distributions are significantly affected by the new network structure. In the trophic network case, the slopes are lower and the distributions have a “fatter tail”, meaning that large extinction events are more common. A microbial community whose interaction network is organised along strongly hierarchical lines is for this reason predicted to be less resilient.

**Fig. 4.**
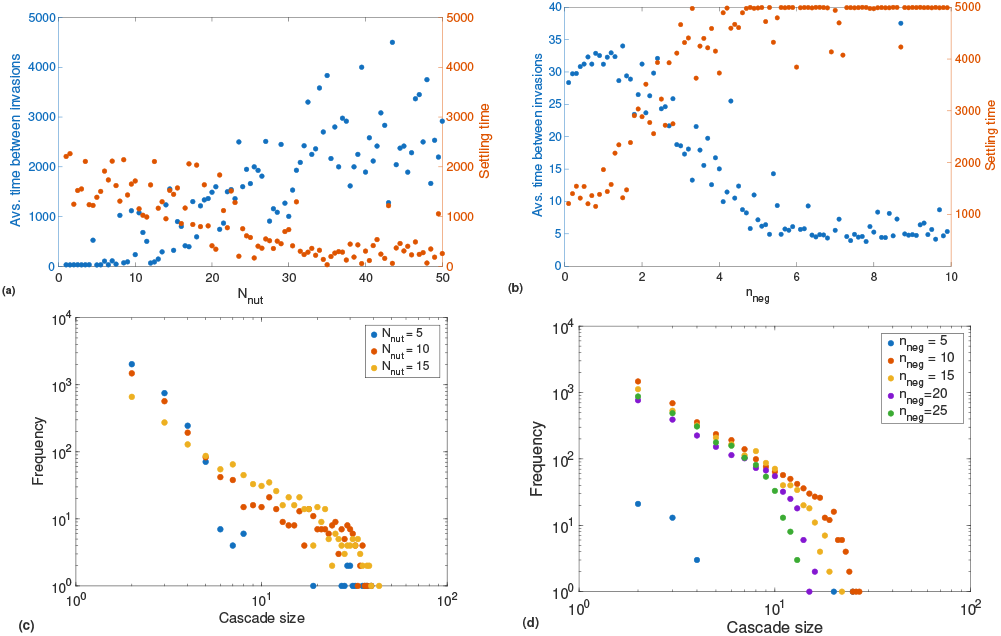
Average time between invasions, settling times (a-b), and cascade sizes (c-d) as functions of the number of nutrients (a,c) and negative links (b,d) for the trophic network model. In comparison with figs. 2 and 3, the dependence of the invasibility measures on *n*_*neg*_ is nearly identical, but stability now increases linearly with *N*_*nut*_. Notably, the cascade size distributions have a lower slope and a fatter tail than in fig. 2, indicating that large extinction events are more likely in the trophic network model. The parameters used are as in fig. 3.

We might expect a network with a skewed degree distribution to result in an unstable community if the numbers of producers and consumers of a particular nutrient are anticorrelated. This is unlikely to be the case in nature, since auxotrophies - the lacking ability of a species to produce an essential amino acid - tend to arise and become more widespread where the “missing” amino acid is widely available in the community (26). On the other hand, if microbial interaction networks are nested in the same way that plant-animal networks are (15), this asymmetry could in principle have a similar effect. The pattern we model of complex nutrients being broken down to simpler ones by a few specialist degraders is another asymmetry that might cause instability. We do in fact see that the tail of the cascade size distribution becomes a lot fatter in the trophic network case. Creating a hierarchical network of nutrient production and consumption by dividing the community into primary, secondary, and higher-level consumers thus appears to enable perturbations in the lower levels to propagate to higher ones.

## Discussion

Despite the simplicity of our model we analytically predict and numerically observe complex dynamics of assembly and collapse under continual invasions by new species. This demonstrates that the model in question is able to capture some of the phenomena that we are interested in studying in real microbial communities. The analytical and numerical results both show that negative interactions between species play a large role in shaping communities. This is true both in regard to their diversity, composition, and stability to invasion and perturbation. The diversity of a community we predict to be inversely proportional to the fraction of possible negative links that actually exist. The stability against both invasion and extinction cascades additionally but not completely, full ecosystem actually maximizes stability. The value of *p*_*neg*_ that results in maximal stability is large enough to ensure that some species are unable to invade, but small enough that established species are unlikely to be displaced. The importance of negative interactions in stabilizing a microbial community has been described by (25), who nevertheless use a different methodology and therefore arrive at a much higher threshold for system destabilization.

As with *n*_*neg*_, the nutrient requirement parameter *n*_*req*_ does appear to have a major effect on resistance looking at the plot in fig. 3 alone. Looking at the system during the simulations, the apparently highly stable states at high *n*_*req*_ turns out to be the system becoming unable to assemble at all when species are highly demanding. The stable states are therefore actually empty systems.

### Alternative network structures

To investigate the effect of different network structures, we generate a hierarchically organised nutrient-bacteria network mimicking the trophic networks common in natural microbial communities (13). Despite the different structure, the change of network has a small effect on the resistance, for which the plots resemble those of fig. 3. One exception is the invasibility as a function of available nutrients *N*_*nut*_, which is much more variable than in the random network case. The resource production parameter *n*_*pro*_ exhibits a similar effect in both network types. Number of required nutrients per species, *n*_*req*_, is not meaningful for the trophic model, since this varies with position in the trophic hierarcy. Plots of the invasibility and settling times as functions of *n*_*neg*_ and *N*_*nut*_, and of the corresponding cascade size distributions are shown in fig. 4 for comparison. On the other hand, the slope of the cascade size distributions are significantly affected by the new network structure. In the trophic network case, the slopes are lower and the distributions have a “fatter tail”, meaning that large extinction events are more common. A microbial community whose interaction network is organised along strongly hierarchical lines is for this reason predicted to be less resilient.

We might expect a network with a skewed degree distribution to result in an unstable community if the numbers of producers and consumers of a particular nutrient are anticorrelated. This is unlikely to be the case in nature, since auxotrophies - the lacking ability of a species to produce an essential amino acid - tend to arise and become more widespread where the “missing” amino acid is widely available in the community (26). On the other hand, if microbial interaction networks are nested in the same way that plant-animal networks are (15), this asymmetry could in principle have a similar effect. The pattern we model of complex nutrients being broken down to simpler ones by a few specialist degraders is another asymmetry that might cause instability. We do in fact see that the tail of the cascade size distribution becomes a lot fatter in the trophic network case. Creating a hierarchical network of nutrient production and consumption by dividing the community into primary, secondary, and higher-level consumers thus appears to enable perturbations in the lower levels to propagate to higher ones.

## Discussion

Despite the simplicity of our model we analytically predict and numerically observe complex dynamics of assembly and collapse under continual invasions by new species. This demonstrates that the model in question is able to capture some of the phenomena that we are interested in studying in real microbial communities. The analytical and numerical results both show that negative interactions between species play a large role in shaping communities. This is true both in regard to their diversity, composition, and stability to invasion and perturbation. The diversity of a community we predict to be inversely proportional to the fraction of possible negative links that actually exist. The stability against both invasion and extinction cascades additionally clearly depend on the number of nutrient requirements per species, with more required nutrients leading to more large cascades but fewer apparent invasions. This latter conclusion may however be due simply to the fact that fewer species can establish themselves in the system overall if their nutrient requirements are very large.

Interestingly, the effect of network structure on the dynamics of this model appears to be modest. Both when using Erdős-Renyi networks and a more structured trophic network, we see that it is mainly the number of negative interactions between species and the nutritional requirements of the average species that determine whether a community can assemble and remain stable. We take the fact that both network structures result in qualitatively similar conclusions as a demonstration that the model is relatively robust to changes in assumptions made about the community and interactions.

In naturally occurring microbial communities, co-occurrence networks seem to have broad degree distributions, often approximating poewrlaws with exponents of between − 1 and − 2 at low connectivities and a steeper cutoff at higher connectivities (22, 27). This pattern is not reproduced by either network model used here. However, such co-occurrence networks might paint over significant complications, especially in the trophic case. For example, if species *A* produces metabolite *a*, which is necessary for species *B* producing metabolite *b*, which then in turn supports the existence of species *C*, we will in a co-occurrence network simply observe coexistence between species *A, B*, and *C*, with no direct information about the trophic hierarchy. Furthermore, while a correlation between two species may be due to a symbiotic relationship between them, it might just as well be due to an underlying interaction between both species and the same nutrient source. Co-occurrences are thus rather evidence of the *lack* of strong antagonism than of symbiosis.

As an additional note, a significant loss of information happens when moving from a consumer-resource graph to a co-occurrence network. For example, consider a perfectly tree-like trophic network, where each producer supports a fixed number *d* of consumers on the trophic level above it. Two trophic levels above, it thereby indirectly supports *d*^2^ species and so on, such that the number of co-occurring species increases exponentially with the number of trophic levels. Taken together, this regular trophic graph results in a scale-free co-occurrence network with exponent − 1. The structure of the co-occurrence network is very different from the consumer-resource one. However, such indirect relationships between species might possibly underpin the observation that such networks are commonly broad-scale. The consumer-resource networks used here do generate a decreasing out-degree distribution, i.e., few species support many others. This is roughly in keeping with what we would expect in a naturally occurring interaction network. For this reason, and due to the doubts about the applicability of co-occurrence networks to this case, we are not too concerned about the deviation of our trophic model network from most of these.

In the present manuscript, we have constructed a simple model of microbial interactions and community assembly, leaving out all but the most basic species-species interactions: exometabolite production and consumption, and antagonistic exclusion of other species.

While not attempting realism, we have nonetheless considered the possible uses of the model in describing real microbial ecosystems. We see two primary potential use cases. The first concerns a situation where a large number of similar communities are well studied, and co-occurrence networks are known to the point where inferences can be made about interspecies interactions. This could for example be a set of soil microbiomes from a homogeneous and well-studied biotope. Alternatively, it could be one of the relatively few nearly completely described animal microbiomes, such as the human nasal microbiome (28, 29) or the bee gut microbiome (30). Here, we could imagine using a model similar to the present one to make predictions about related but as-yet unobserved communities and to inform the inferences from species correlations to causal, interactional links between species. This method would require an assumption that the studied community is stable, at least on short or intermediate timescales.

The other case is the converse situation, where the set of species likely to be found in the environment are well-studied, and interactions between them well established, but where the full makeup of the community is less clear. An example of this could be the main taxa of bacteria likely to be found in the human gut, whose interaction parameters are somewhat quantifiable (31), but whose interactions in an ensemble are poorly understood. In this case, the model might help predict the final makeup, and especially the diversity and stability of a community based on the available species.

It would greatly improve the applicability of this model, and might lead to new insights in microbiome dynamics if one tested its predictions in ecosystems where eqs. (2-6) predict instability or small diversity. Eq. (2) predicts that microbial ecosystems with many antagonistic interactions have low diversity. Communities that have long co-evolved in nutrient-poor environments, and which have therefore emphasized competitive interactions might provide a natural example of this.

As for stability and resilience, eqs. 4 and 6 describe two conditions for this, namely that the number of nutrients involved in the community metabolism should be smaller than the number of species, and the expected stable number of resident species, 1*/p*_*neg*_, should not be much smaller than the number of species present in the environment. We would imagine that, e.g., a small gut microbiome exposed to an extremely diverse diet would violate the first condition, and that a small isolated community exposed to a new, more open environment would violate the second. We predict that under such circumstances, one would observe a greatly decreased stability of the community. Notably, both suggested examples deal with systems experiencing a perturbation, since we do not believe that such conditions would be stable over time.

In this article, we have presented a new, simple model describing the assembly of microbial communities, and have thereby derived conditions for their stability over time. Though we have outlined a few cases when our model might have predictive power, we have so far refrained from attempting to reconstruct any real communities using this model, and from trying to parameterize it using data from such communities. The reason for this omission is the vast diversity of relevant communities and a desire to present a framework for understanding them that, while abstract, is also general. It is our hope that researchers more intimately familiar with specific microbial communities of the soil, the ocean, or animal microbiomes may find this framework useful and let themselves be inspired to create similar models appropriate for their area of the microbial world. Thereby, we believe that this theoretical framework can help better understand the rules governing community assembly.

## Data, Materials, and Software Availability

The code needed to generate the figures shown here can be found on Figshare (DOI: 10.6084/m9.figshare.30898922).

## ACKNOWLEDGMENTS

We would like to thank Jordi Bascompte for his encouragement and many useful comments on the manuscript. Furthermore, we thank Alberto Pascual and Sara Mitri for enlightening discussion and suggestions.

