## Supplementary material for "Modeling assembly dynamics and stability of microbial communities": SI Appendix

### Modeling assembly dynamics and stability of microbial communities - supplementary material

December 2025

This supplementary material contains the numerical verification of the analytical results derived in the article "Modeling assembly dynamics and stability of microbial communities". In addition, it contains plots of the results obtained using the trophically inspired network where these are not interestingly different from the random network results, as well as a closer look at the temporal dynamics of the ecosystem in some specific cases.

Here follows a summary of the parameters used:

- $p_{neg}$: the probability of a negative link existing between any species $i$ and any other species $j$, i.e., that $j$ cannot exist if $i$ is present.
- $p_{pro}$: the probability that any species $i$ produces any nutrient $k$.
- $p_{req}$: the probability that any species $i$ requires any nutrient $k$ to invade or remain in the system.
- $N_{nut}$: the number of nutrients used or produced by any member of the community.
- $N$: the maximum number of species potentially present in the community. Can be taken to represent the number of species available in the greater environment.
- $n_{neg}, n_{pro}, n_{req}$: the mean numbers of outgoing or incoming negative links, nutrients produced, and nutrients required per species. Related to the corresponding probabilities as $n_{neg} = p_{neg}N$, $n_{pro} = p_{pro}N_{nut}$, $n_{req} = p_{req}N_{nut}$.

#### Verification of the analytical results

Using the criteria that the system should be resistant to invasion, or alternatively at a dynamic equilibrium wrt. diversity (defined as number of resident species), we derive the equilibrium diversity $x^*$:

$$x^* = 1/p_{neg}. \tag{1}$$

We test this numerically by running our model, keeping all parameters constant except $p_{neg}$. The graph of the resulting diversities can be seen in fig. 1. Changing the other parameters by a factor of around two has little effect. Only when reducing the total number of species $N$ do we see that the function $x^*(p_{neg})$ becomes flatter. $x^*$ is analytically predicted to be independent of $N$, but of course there can be no more resident species than there are species available in the environment.

We see that the numerical results behave as expected, except at very low $p_{neg}$ where the diversity is lower in the numerical model. One possible reason for this is the sorting effect brought up in the main article as well: highly competitive species that themselves are difficult to outcompete are likely to become established first, meaning that the resident species at any given time will have an above-chance number of negative out-links and a below-chance number of negative in-links. This effect is not accounted for in the analysis.

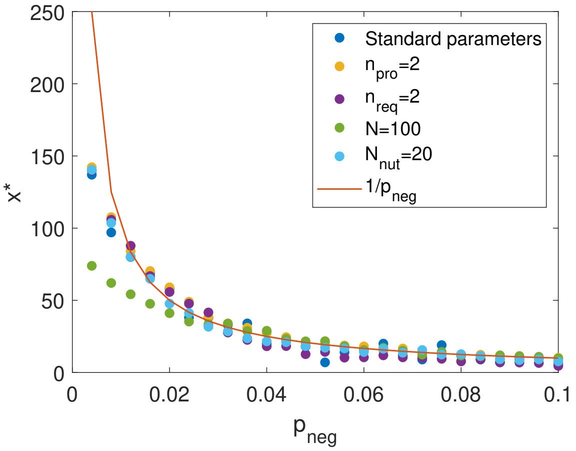


Figure 1: Numerical measurement of $x^*$ (measured at the end of the simulation) in the model vs. the analytical result $x^* = 1/p_{neg}$. The other parameters are $N = 250$, $N_{nut} = 10, n_{pro} = 1, n_{req} = 1$.

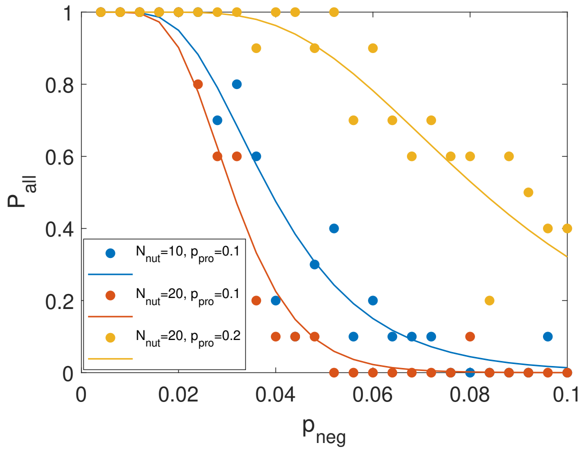


Figure 2: Numerical (dots) and analytical (lines) results for the probability that all nutrients are produced at equilibrium, $P_{all}$. Here, $n_{req} = p_{req}N_{nut} = 1$.

Similarly, we measure the probability $P_{all}$ of all nutrients being produced at equilibrium in the simulation. The measurement is carried out after 2500 timesteps. As derived in the main article, this should be given by

$$P_{all} = [1 - (1 - p_{pro})^x]^{N_{nut}} = [1 - (1 - p_{pro})^{1/p_{neg}}]^{N_{nut}} \tag{2}$$

In fig. 2, we see that the results fit the analytically derived graphs well. To make sure that the dependence on all three parameters, $p_{neg}$, $p_{prod}$, and $N_{nut}$, is as expected we use the former as plotting variable and vary the latter two. Some stochastic fluctuations are to be expected due to the randomness of the networks and invasion dynamics, and since we only run the simulation 10 times per parameter set in the interest of computing time.

Finally, we measure the probability that there are no potential invaders among non-resident species. The result can be seen in fig. 3. We see that the theory predicts a swift transition from nearly no probability of invaders to near-certainty of this at $n_{neg} \approx 1$, meaning $p_{neg} \approx 1/N$. The data do confirm that this is the point at which invaders become more likely, but the transition is a lot more gradual than predicted. The numerical results also do show that the transition shifts to higher $p_{neg}$ at lower $N$, but the slope changes as

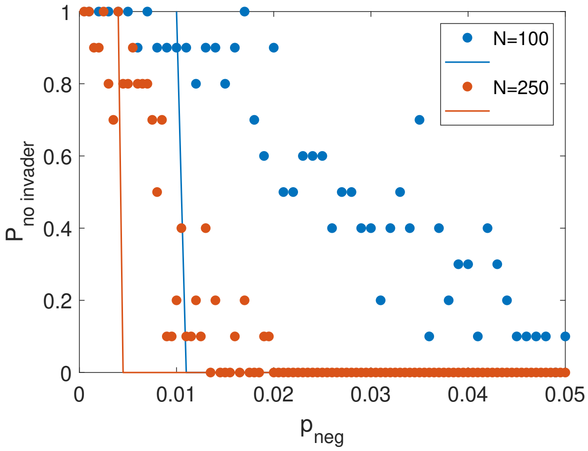


Figure 3: Probability of no potential invaders existing as a function of $p_{neg}$, measured numerically (dots) and calculated analytically (lines). Parameters are as in the previous figures, except for $N$.

well, becoming flatter, which is not captured by the expression. The result is thus only approximate, and we again ascribe the deviation from the numerical result to our analytical assumption that both residents and non-residents are randomly chosen.

### Trophic network - results for varying $n_{pro}$

For reasons of space, and because the results are not interestingly different from the random network model, the plots shown in fig. 4 were left out of the main article. They show the resistance and resilience of the trophic network model as a function of $n_{pro}$, as measured using time between invasions, settling time (that is, time until last successful invasion, after which the system is "settled down"), and cascade size distribution. We only vary $n_{pro}$ up to five, since for larger numbers, each species will produce such a large fraction of the nutrients ($N_{nut} = 10$ as in the main article) that it is difficult to maintain the hierarchical structure of the network.

### Dynamics behind invasibility plots

In this section, we display some of the invasion-extinction dynamics that may not be conveyed by the aggregated data of the main article. The plots of figs. 5-8 show species existence (yellow pixels) or non-existence (blue pixels) as a function of time for various parameter sets. We have aimed to include plots of as diverse a set of parameter regimes as possible. The standard parameter set unless otherwise indicated is $N_{nut}$=10, $n_{neg} = 1$, $n_{pro} = 1$, $n_{req} = 1$, and $N = 250$.

It can be seen that for all cases where $n_{neg}$ is small ($n \lesssim 1$), the system sooner or later settles down to a stable, non-invasible state. Only when $n_{neg}$ is significantly larger than 1 does the system remain unstable throughout the simulation. The other parameters mainly seem to affect the settling time. A single exception is $n_{req}$, which prevents a community from assembling at all when it is too high, representing all available species being too picky for the set of nutrients potentially present in this ecosystem. These snapshots of the dynamics serve to show the actual system states underlying the stability graphs of the main article, and show among other things that the system for high $n_{req}$ is not in fact an extremely stable community, but simply the absence of a community, while other instances of stability do look as we would expect.

Finally, in fig. 9 we showcase a couple of examples of the dynamics of the trophic network model, illustrating the larger extinction cascades of this version. Also notable is the fact that even for relatively low $n_{neg}$, the system may have difficulties assembling. The more refined nutrients high in the trophic hierarchy

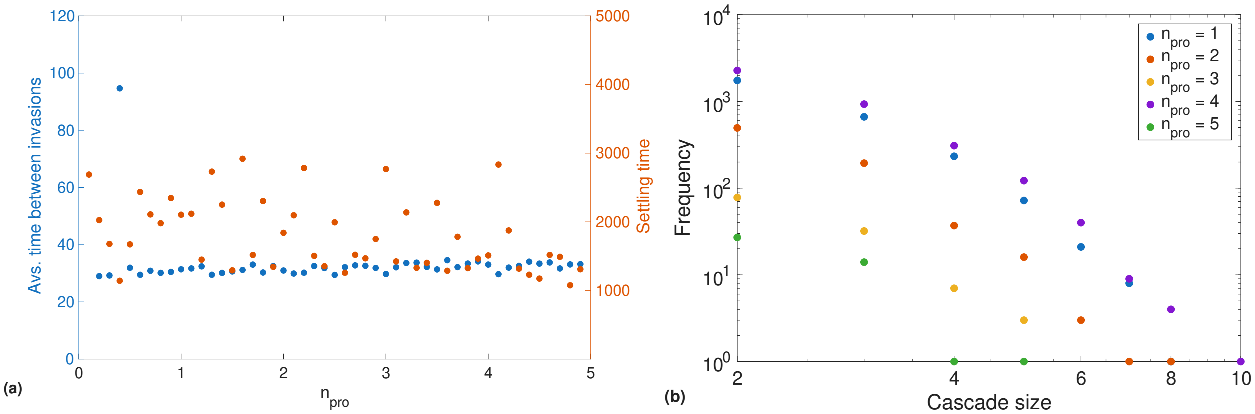


Figure 4: Results of simulations using a trophically inspired network nutrient network structure. (a) The two measures of invasibility, time between invasions and settling time (time to last successful invasion). Note that due to outliers, the left y-axis is different from the corresponding non-trophic figure in the main article, which is the main reason the two figures look different. (b) Frequency of extinction cascades as a function of their size. Parameters: $N = 200$, $N_{nut} = 10$, $n_{neg} = 1$ in (a) and 5 in (b).

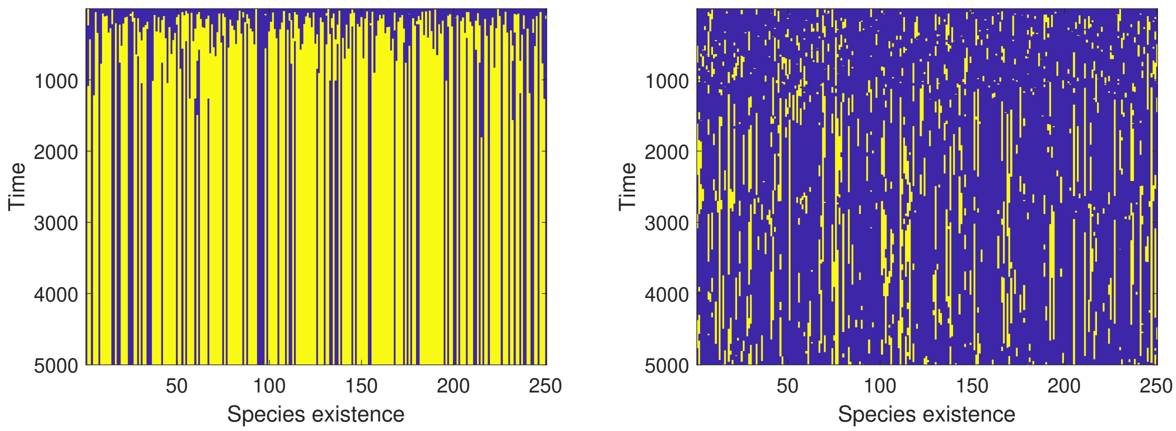


Figure 5: Species existence over time for $n_{neg} = 0.5$ (left) and $n_{neg} = 8$ (right).

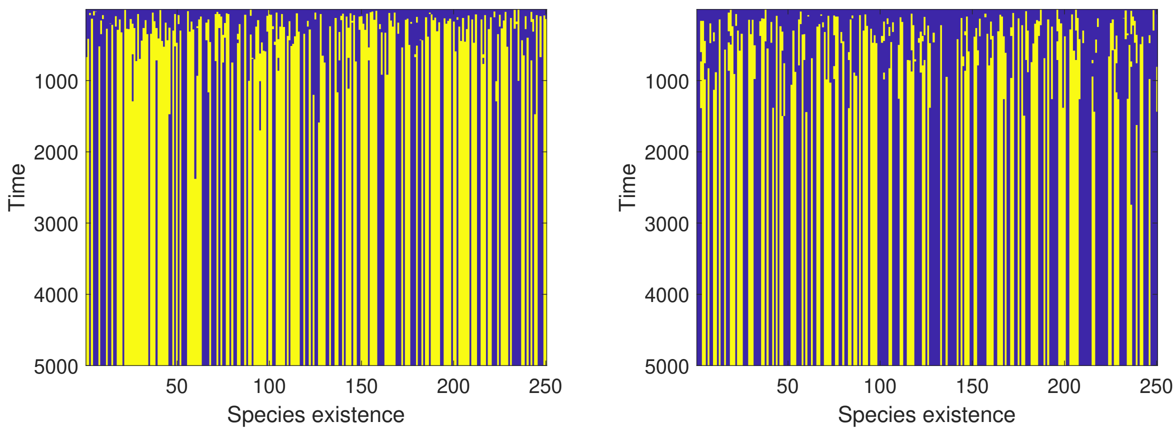


Figure 6: Species existence over time for $N_{nut} = 20$ (left) and $N_{nut} = 80$ (right).

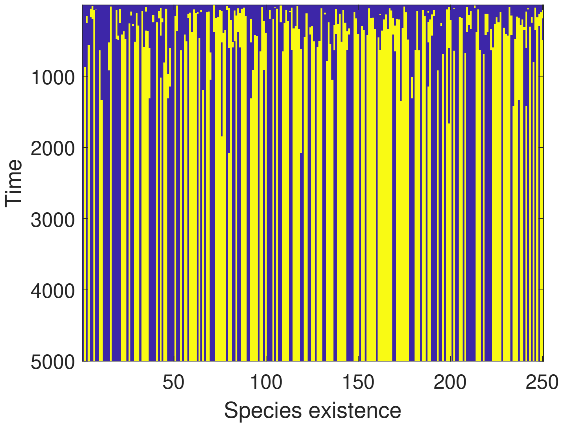


Figure 7: Species existence over time for $n_{req} = 2$ (left) and $N_{req} = 8$ (right). The turquoise field indicates a permanently empty ecosystem.

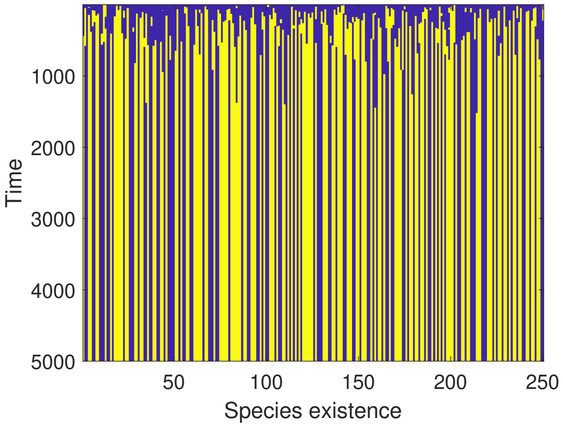


Figure 8: Species existence over time for $n_{pro} = 1$ (left) and $n_{pro} = 8$ (right).

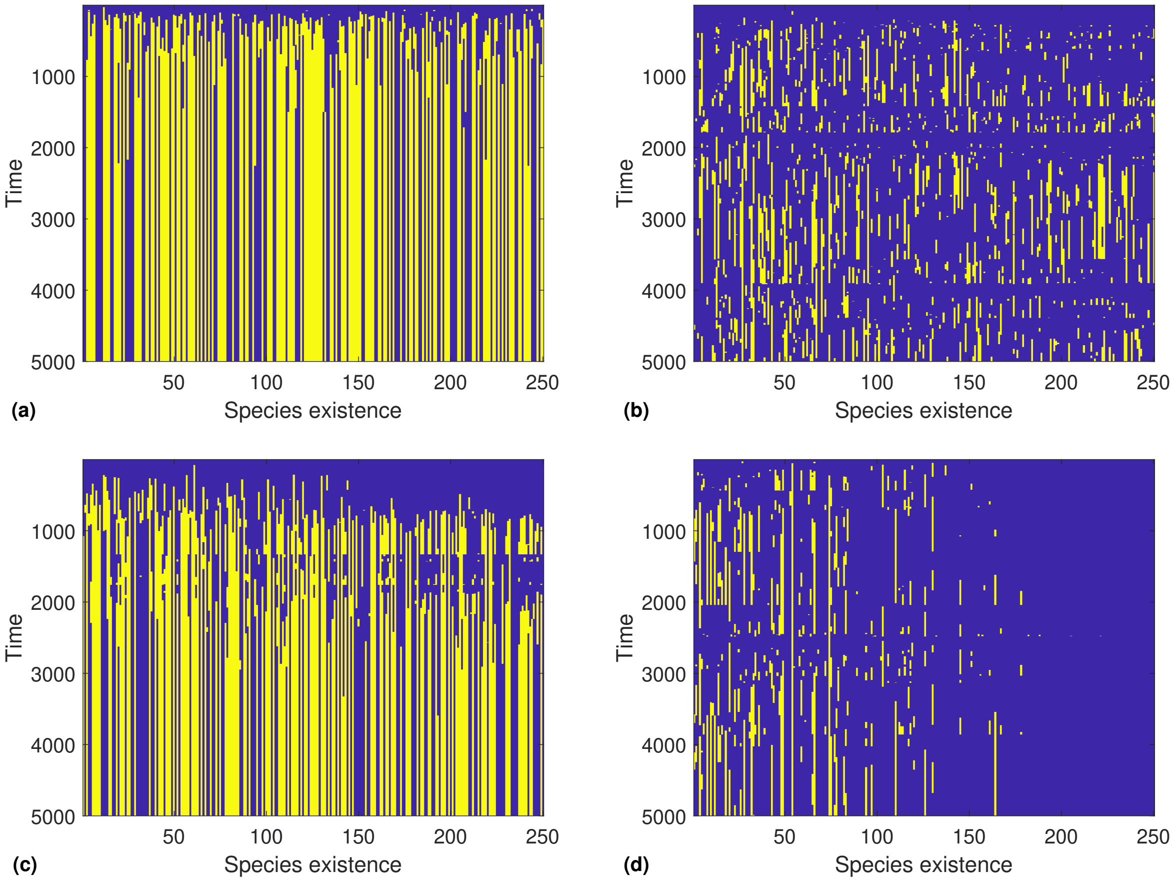


Figure 9: Species existence over time for the trophic network model. (a) Standard parameters, (b)$n_{neg} = 5$, (c) $N_{nut} = 20$, (d) $N_{nut} = 20$ and $n_{neg} = 5$. Extinction cascades can be seen as the yellow lines ending simultaneously. This gives rise to horizontal blue “bands”, indicating a mostly empty community.

seemingly do not get produced, leading to a lower $x$ than would be expected in the random-network case. A plot of $x$ as a function of $p_{neg}$ in the trophic model is shown in fig. 10. The diversity is indeed lower than predicted, but only by a small amount.

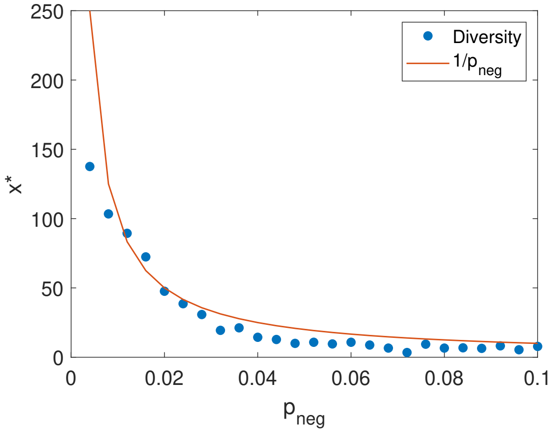


Figure 10: Species diversity $x^*$ at the end of the simulation as a function of $p_{neg}$ in the trophic model. All other parameter values are the standard ones.
